## Supplementary Methods for "When cheating turns into a stabilizing mechanism of mutualistic networks"

### S1, Supplementary Materials for

**This PDF file includes**

### Supplementary methods

*Building initial interaction matrix*

To get a binary interaction matrix, that gathers pairwise ability of species to interact mutualistically, we used trait matching approach. Each species was described by a unidimensional and gaussian morphological trait, $\mathcal{N}(\mu, \sigma)$, with $\mu$ drawn in a uniform distribution $U$(-2.5,2.5) and $\sigma$ in $U$(0.1,0.7). The strength of legitimate interaction ($I_{M_{ij}}$) between a plant *i* and a pollinator *j*, was calculated as the overlap of the densities of the gaussian distribution describing the morphological trait of both species. Thus, $0 < I_{M_{ij}} \leq 1$. To transform this matrix into a binary interaction matrix with a given connectance ($\varphi$), we set all values greater than the quantile 1- $\varphi$ to one, and all values lower than that to zero. Then, we checked that all species had at least one mutualistic partner, and if not, we added one interaction corresponding to the highest value of $I_{M_{ij}}$ relative to that species. The addition of these interactions affected connectance in a negligible way but allowed all species to have at least one mutualistic partner.

*Species growth rates*

Growth rates which were drawn randomly from a beta distribution ($B$), to sample a wide variety of possible vectors of growth rates. We fixed the first shape parameter at $a = 1$ and we drew the second shape parameter $b$ from $e^{U(\log(0.6), \log(3))}$ to sample values between 0.6 and 3, but with more $b$ values around 0.6 than around 3, allowing to model a wide diversity of growth rate distributions. Since we modeled obligatory mutualism in which persistence of species depended on the balance between growth rate and mutualism benefits, to avoid that communities collapsed too often, we bounded the space of growth rates between -0.5 and -0.001:

$$r_{P_i} \text{ and } r_{A_j} \sim min\left(\frac{-1}{2} \times B\left(a = 1, b \sim e^{U(\log(0.7), \log(3))}\right), -0.001\right) \qquad (S1)$$

*Functional responses, and construction of vectors $\Pi$, $\Theta$ and $\Upsilon$*

In our model we included competition for mutualistic partners as interference in the functional response. We did so to account for the fact that illegitimate interactions also affect competition interactions. For each species, competition for mutualistic partners depended on a vector of pairwise competition coefficient, $\Pi$ for plants and $\Theta$ for pollinators. These competition coefficients varied from 0 (no competition) to 1 (maximal competition) and increase if species share common mutualistic partners. The calculation of these vectors can appear unnecessary complex but has for purpose to account for specie in competition strength. Most of the studies model competition strength proportional to the proportion of shared partner, without accounting for the abundance of these partners, meaning for example that even if plants shared by two pollinators become extinct during the simulation, pollinators still compete for them. While our way to calculate competition vectors account for abundance of partners: if a plant species becomes extinct, pollinators stop to compete for it, and conversely. Pairwise competition between a plant *i* and *k*, or between a pollinator *j* and *k*, were calculated as follow:

$$\Pi_{ik} = \frac{1}{\sum_{j=1}^{n_A} M_{ij} \times A_j} \times \sum_{j=1}^{n_A} A_j \times M_{ij} \times M_{ik} \qquad (S2)$$

$$\Theta_{jk} = \frac{1}{\sum_{i=1}^{n_P} \left(M_{ij} + C_{ij}\right) P_i} \times \sum_{i=1}^{n_P} P_i \times \left(M_{ij} + C_{ij}\right) \times \left(M_{ik} + C_{ik}\right) \qquad (S3)$$

For the costs of illegitimate interactions for plants, we used a similar functional response than for mutualistic interactions, with few subtilities. In this case, the interference term was not a competition term anymore but represents the fact that if a cheater interacts illegitimately with many plants, costs of its cheating is diluted among the plants. However, the dilution vector $\Upsilon$ is calculated in an analogous way than vectors $\Pi$ and $\Theta$:

$$\Upsilon_{ik} = \frac{1}{\sum_{j=1}^{n_A} C_{ij} \times A_j} \times \sum_{j=1}^{n_A} A_j \times C_{ij} \times C_{ik} \qquad (S4)$$

*Setting values for functional response parameters*

To avoid that system collapse, species per-capita benefits from mutualistic interaction should buffer mortality and intra specific competition. For example, for pollinators, that means that $r_{A_j} + \frac{\sum_{i=1}^{n_P}[\alpha(M_{ij}+C_{ij})-\Lambda M_{ij}]P_i}{1+\beta\sum_{i=1}^{n_P}(M_{ij}+C_{ij})P_i+c\sum_{k=1}^{n_A}\Theta_{jk}\times A_k} - \sum_{k=1}^{n_A} c_{A_{jk}} \times A_k = 0$ has a solution with $A_j > 0$. To do so, per-interaction benefit of mutualism should be high enough to allow species to persist in theory when there is no cheating, but not too high otherwise persistence would be always maximal and never vary. In a case without cheating, when the summed abundance of mutualistic partners tends to infinity, the functional response tends to $\frac{\alpha-\Lambda}{\beta+c}$ so towards $(\alpha - \Lambda)/2$, because we set $\beta = 1$ and $c = 1$. Since growth rate can reach -0.5 and $\Lambda = 0.3$, we chose $\alpha = 1.5$, which is close to the minimal per-interaction benefits that allow species to persist even in those conditions.

*Two scenarios of simulations: cheaters are the generalist pollinators or the specialist ones*

Since we expected different outcomes of cheating depending on the fact that the cheaters were the specialist (with few partners) or the generalist (with many partners) pollinators, we performed simulations for these two cases. To quantify the specialism/generalism degree of pollinator $j$, we estimated the diversity of its interaction partners, using the Shannon index over $I_{M.j}$. Then, in the case in which cheaters were specialist species, we defined the $round(\overline{\Delta} n_A)$ species with the lowest Shannon index as cheaters. In the case in which cheaters were generalist species, we defined the $round(\overline{\Delta} n_A)$ pollinator species with the highest Shannon index and with maximum $n_A - 1$ mutualistic partners as cheaters. That means that if a pollinator was perfectly generalist, so interact with every plant species, it was not defined as a cheater, because we wanted to study cases in which cheating can be conservative (within the mutualistic niche) or innovative (outside the mutualistic niche). If the cheater is perfectly generalist, then innovative cheating is not possible anymore.

*Measuring plant traits*

While putting camera trap on the field, flowers were collected, and few traits were measured on standardized pictures. The length of the corolla, of the stigma and anther and the curvature of the corolla. To measure curvature, an imaginary circle was fitted to the corolla shape and the curvature was measured as one divided by the radius of the circle.

*Additional references*

Nakajima, H. & Higashi, M. (1995). Indirect effects in ecological interaction networks II. The conjugate variable approach. *Math. Biosci.*, 130, 129–150.
