## Supplementary Figures and Tables for "When cheating turns into a stabilizing mechanism of mutualistic networks"

### S2, Supplementary Materials for

**This PDF file includes**

Supplementary figures........................................................................................................ 2
Supplementary tables ........................................................................................................ 10

### Supplementary figures

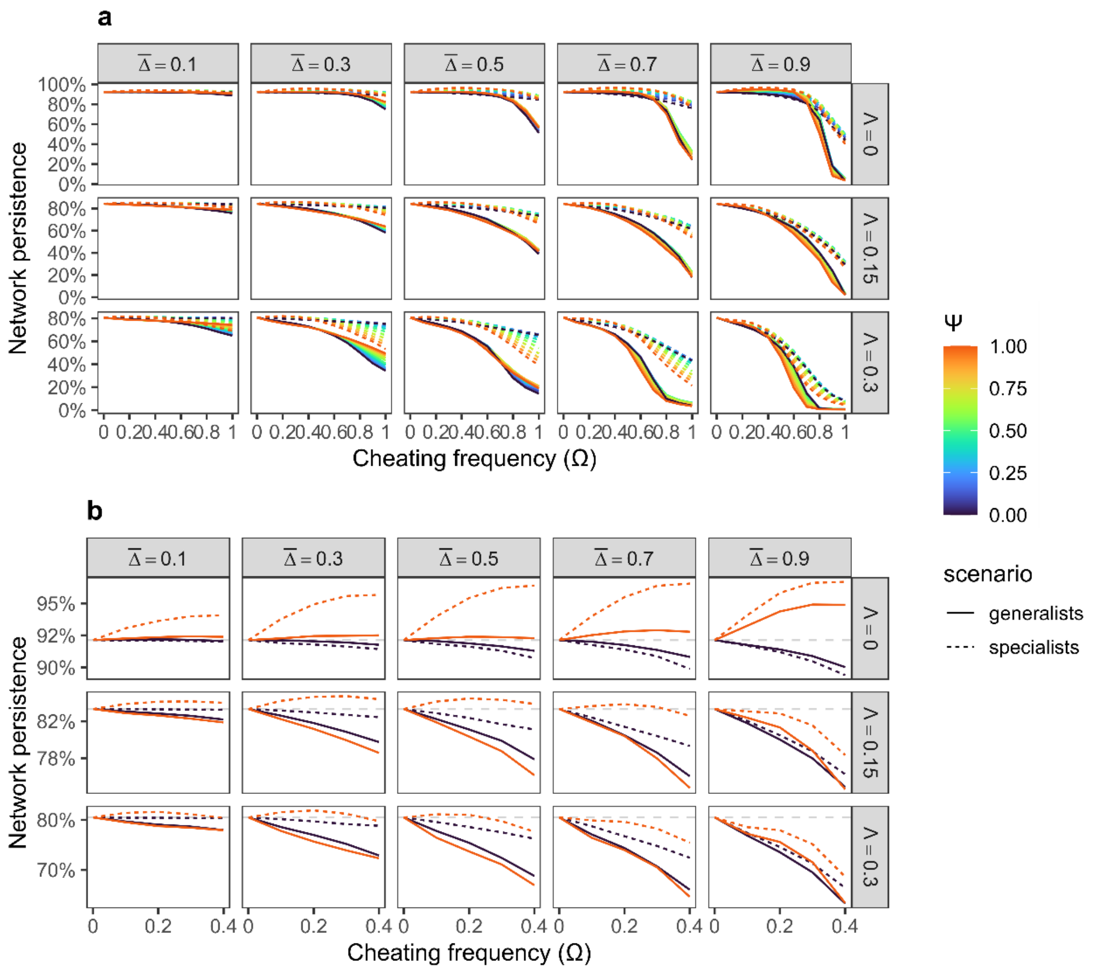


***Figure S1: Innovative cheating can increase network persistence.*** *(a) Network persistence as a function of cheating frequency (Ω), proportion of cheaters ($\bar{\Delta}$), cost associated with mutualism (Λ) and proportion of innovative cheating (Ψ). Network persistence is measured as percentage of species (plants and pollinators) having positive abundance at equilibrium, relative to the total initial number of species. Dashed lines show average results when cheaters are the most specialized species and solid lines when cheaters are the most generalist species. Here results are presented for initial values of connectance φ=0.4 and averaged over the 500 initial conditions. (b) Shows the same than in (a) but on a truncated axis of cheating frequency and for two extremes values only, when cheating is conservative only (Ψ=0) or innovative only (Ψ=1). The dashed grey line shows the average network persistence when there was no cheating. Here results are presented for initial values of connectance φ=0.4 and averaged over the 500 initial conditions. A cost of Λ=0.15 and Λ=0.3, corresponds to 10% and 20% of benefits associated with mutualism, respectively (Λ/α = 0.1 and 0.2).*

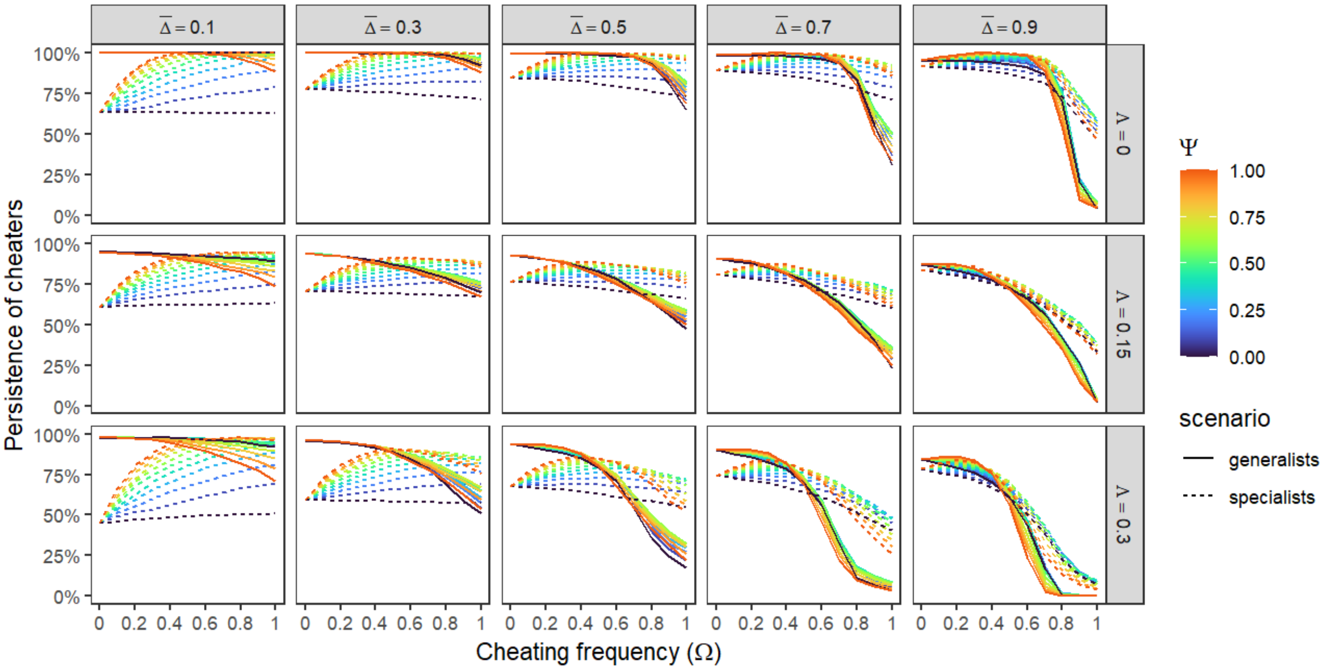


***Figure S2: Effects of cheating on cheater persistence depend on the generalism degree of the cheaters.*** *Cheater persistence (i.e. number of initial cheaters divided by number of cheater at equilibrium) as a function of cheating frequency (Ω), proportion of cheaters ($\bar{\Delta}$), cost associated with mutualism (Λ) and proportion of innovative cheating (Ψ). Dashed lines show average results when cheaters are the most specialized species and solid lines when cheaters are the most generalist species. Here results are presented for initial values of connectance φ=0.4 and averaged over the 500 initial conditions. We can see that for low and intermediate level of cheating, innovative cheating tends to increase the persistence of cheaters, if they are specialists, while it is the opposite if they are generalist.*

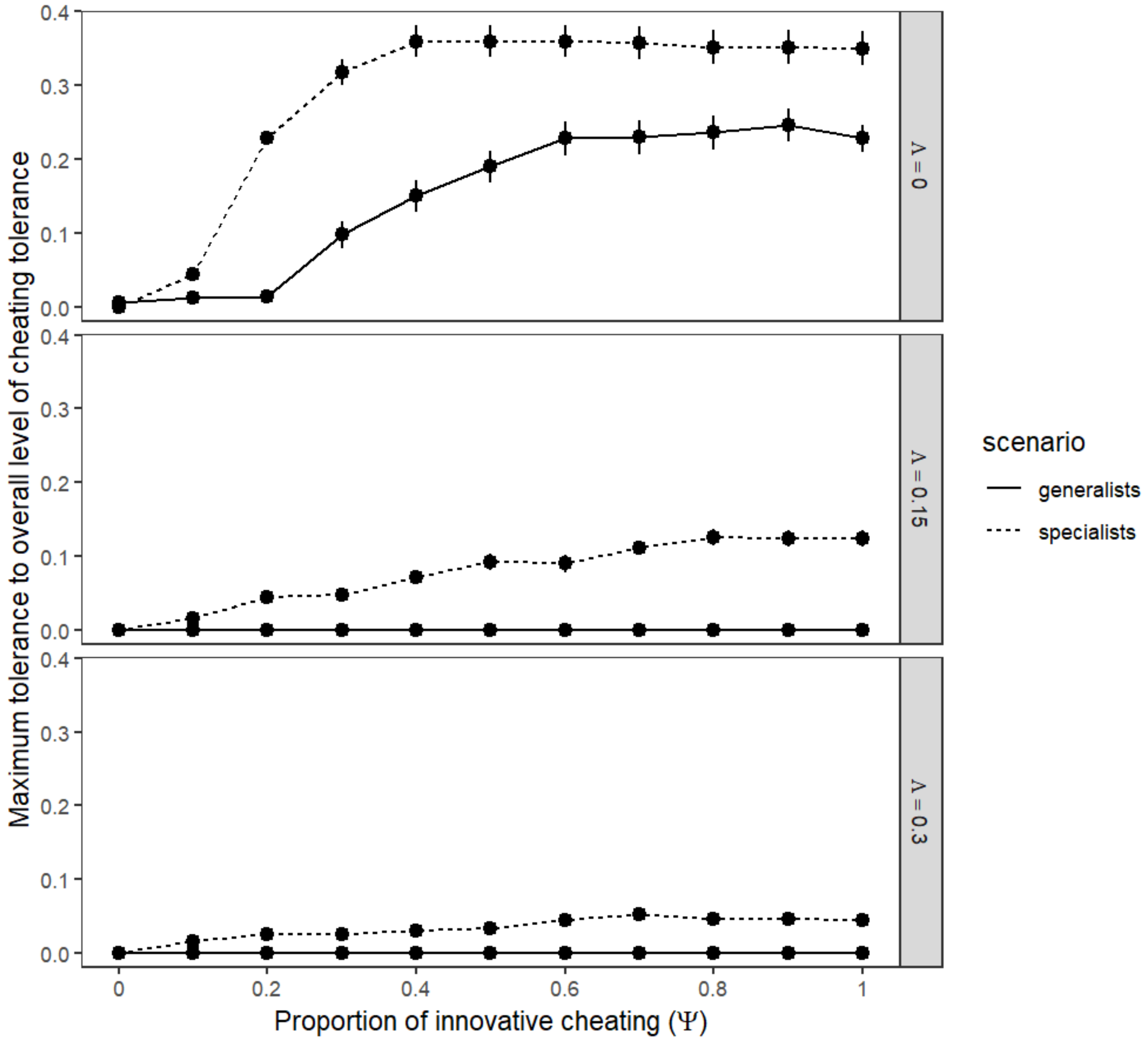


***Figure S3: Maximum overall level of cheating allowing positive effects on persistence increases when cheating is innovative but decreases with cost associated with mutualism.*** *Tolerance to cheating, calculated as the maximal value of $\bar{\Delta} \times \Omega$ leading to positive effects on network persistence in average, as a function of proportion of innovative cheating (Ψ), cost associated with mutualism (Λ) and the scenario of simulations, cheaters are generalist species or specialist species. Here results are presented for initial values of connectance φ=0.4, and results are averaged over the 500 initial conditions and over all values of $\bar{\Delta}$ and Ω. Error bars are the standard errors associated with average values.*

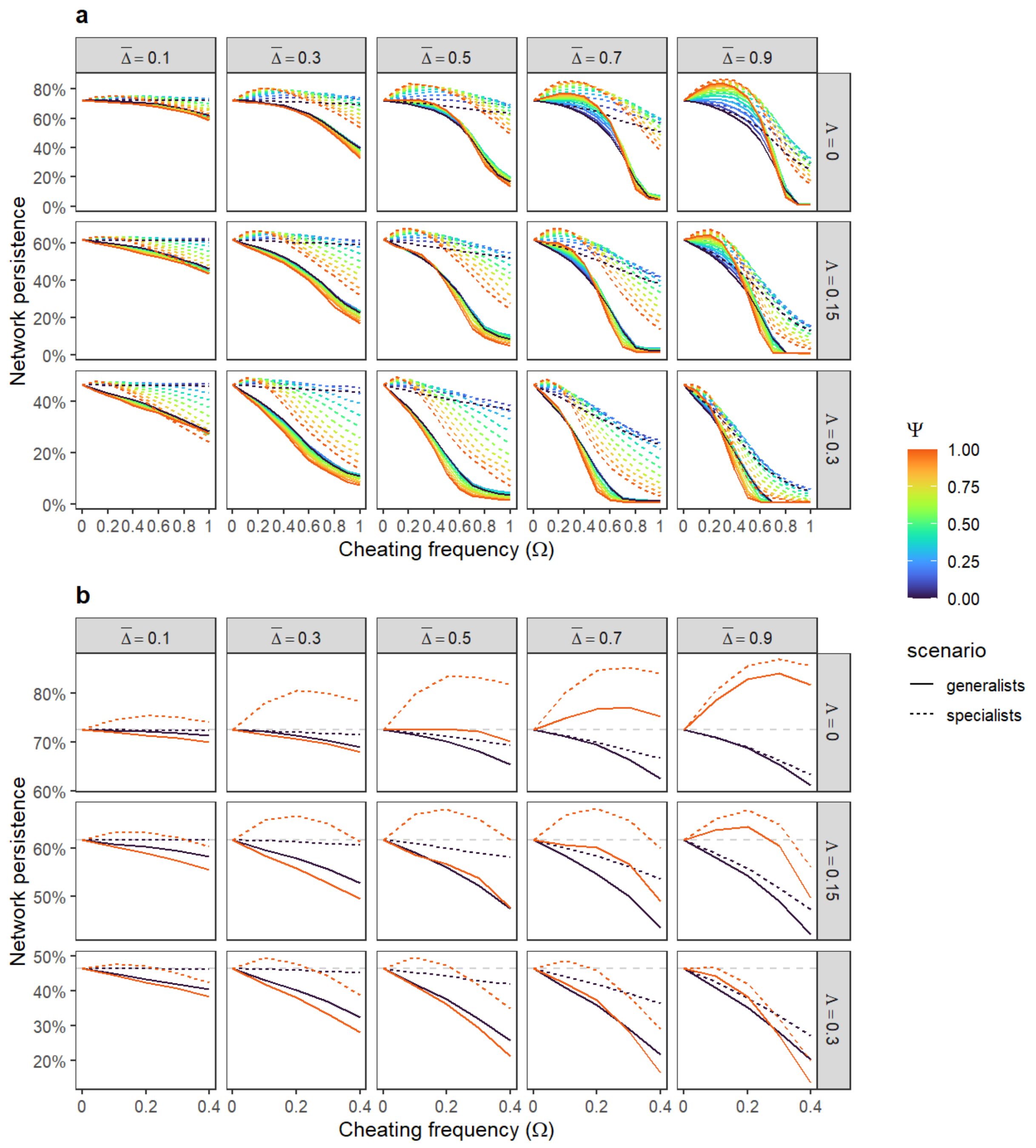


***Figure S4: Same figure than Fig. S1 but with a lower value of initial connectance ($\varphi$=0.2).*** *(a) Network persistence as a function of cheating frequency ($\Omega$), proportion of cheaters ($\bar{\Delta}$), cost associated with mutualism ($\Lambda$) and proportion of innovative cheating ($\Psi$). Dashed lines show average results when cheaters are the most specialized species and solid lines when cheaters are the most generalist species. Here results are presented for initial values of connectance $\varphi$=0.2 and averaged over the 500 initial conditions. (b) Shows the same than in (a) but on a truncated axis of cheating frequency and for two extremes values only, when cheating is conservative only ($\Psi$=0) or innovative only ($\Psi$=1). In (b) the dashed grey line shows the average network persistence when there was no cheating.*

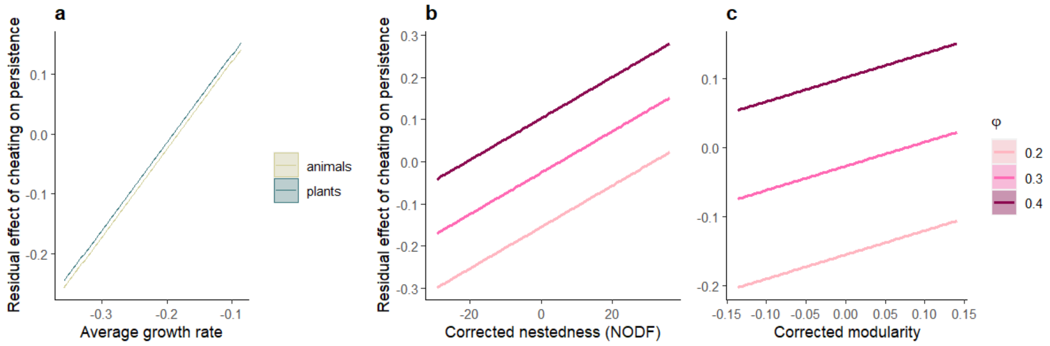


***Figure S5: Effect of cheating on persistence is linked to plant-pollinator dependency and to network structure.*** *Residual effects of cheating on persistence as a function of (a) average plants and animals' growth rates, (b) connectance and nestedness, and (c) connectance and modularity of the initial mutualistic network (*$I_M$*, cf. Methods). Residual effects of cheating are calculated by subtracting the average effect corresponding to the respective parameter combination from the effect observed for each individual simulation. Positive values indicate effect of cheating on persistence above average effect while negative values indicate the opposite. In (a), more negative growth rates indicate stronger dependency to the other guild. Lines represent model predictions. Nestedness and modularity measures were corrected by connectance (*$\varphi$*) to disentangle their respective effects.*

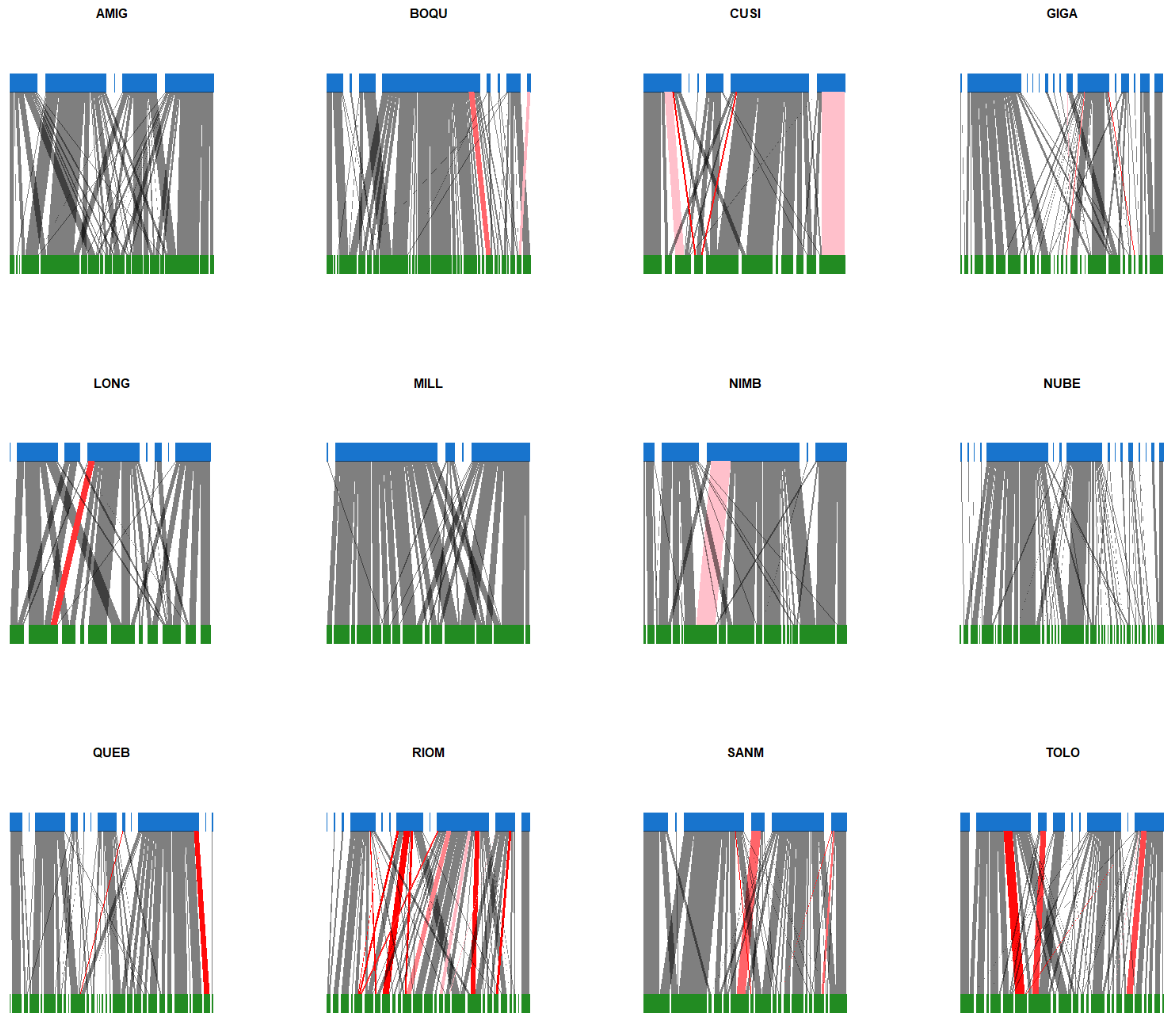


***Figure S6: Plant-hummingbird interaction network from Costa Rica.*** *Empirical network of interactions between plants (green) and hummingbirds (blue) from Costa Rica for each site. Thickness of the lines is proportional to interaction frequency, while colour of the lines represents cheating frequency, grey (only legitimate) and from pink (low cheating frequency) to bright red (only illegitimate).*

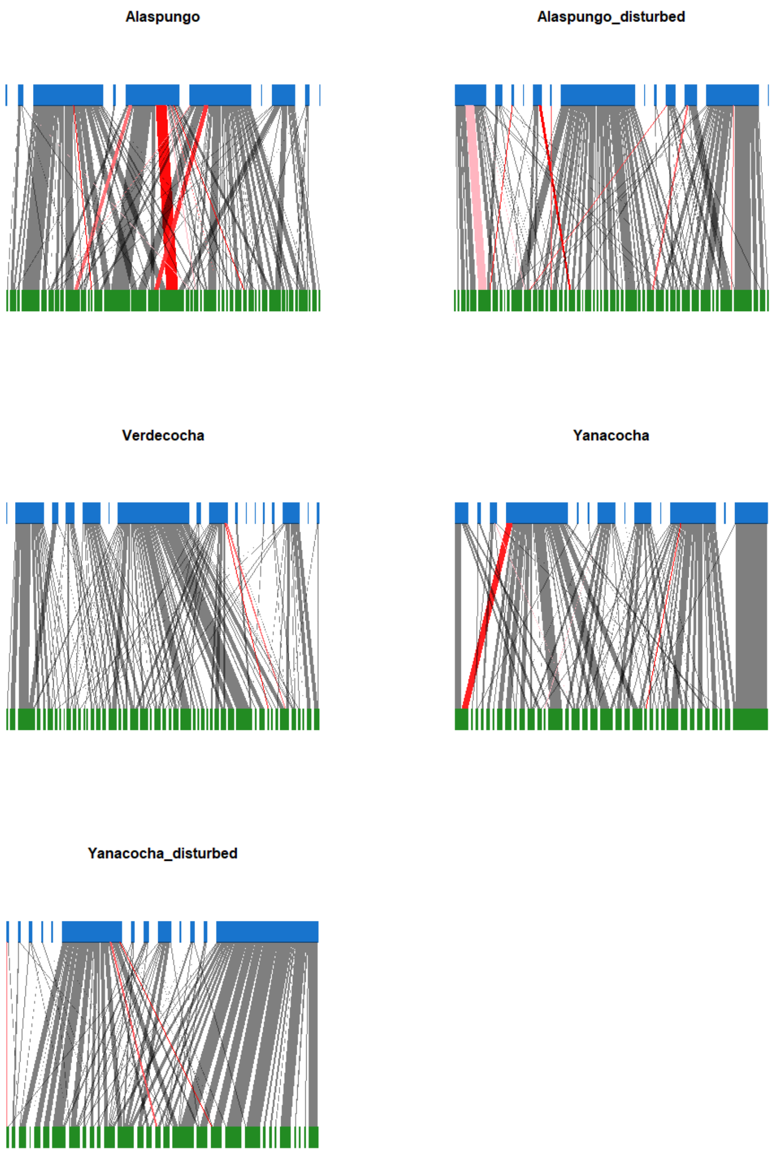


***Figure S7: Plant-bird interaction network from Ecuador.*** *Empirical network of interactions between plants (green) and birds (blue) from Ecuador for each site. Thickness of the lines is proportional to interaction frequency, while colour of the lines represents cheating frequency, grey (only legitimate) and from pink (low cheating frequency) to bright red (only illegitimate).*

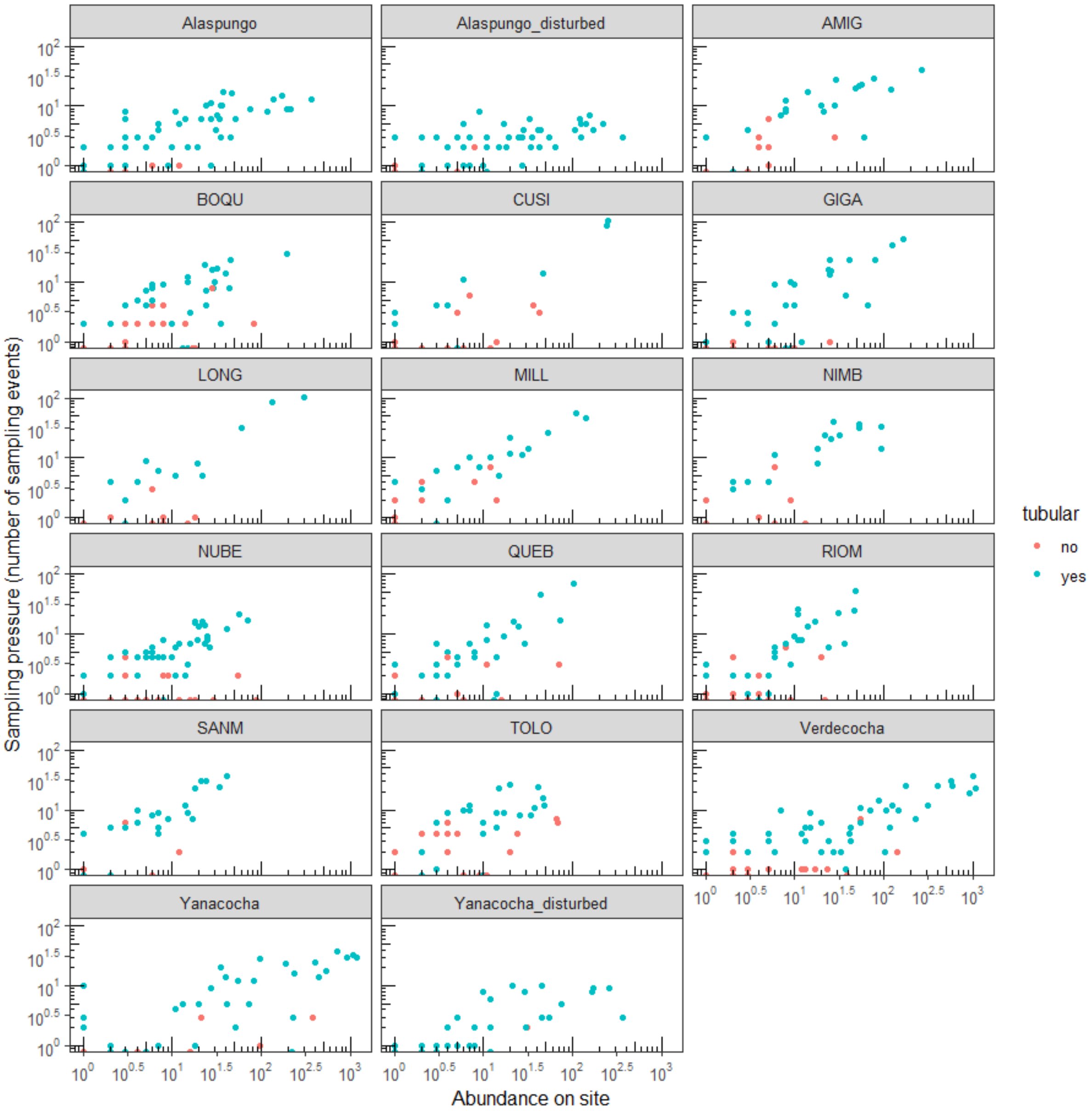


***Figure S8: Sampling pressure as a function of plant abundance, for each site.*** *Sampling pressure, the number of sampling events done on one species on a given site, as a function of the abundance of flowering plants on that site. Each point is a plant species. Colors indicate if there is, or not, a closed corolla forming something roughly similar to a tube. Tubular flowers are more likely to be visited by hummingbirds.*

#### Supplementary tables

**Table S1:** Results of type II ANOVA of the three models presented in Fig. 4b,d,e. Df = degree of freedom.

| Response variable | Explicative variable | $\chi^2$ | Df | p-value | Marginal $R^2$ |
|---|---|---|---|---|---|
| Frequency of cheating | Diversity (log) | 4.2872 | 1 | 0.03840 | 0.11885 |
| | Country | 0.0001 | 1 | 0.99154 | |
| | Country:Diversity (log) | 0.2740 | 1 | 0.60066 | |
| Proportion of cheaters | Elevation | 4.6810 | 1 | 0.03050 | 0.28708 |
| | Country | 3.3014 | 1 | 0.06922 | |
| | Country:Elevation | 0.1545 | 1 | 0.69423 | |
| Overall level of cheating | Elevation | 8.9353 | 1 | 0.00280 | 0.46702 |
| | Country | 3.1084 | 1 | 0.07789 | |
| | Country:Elevation | 0.0047 | 1 | 0.94531 | |

**Table S2:** Locations of the studied sites.

| Site | Longitude_WGS84 | Latitude_WGS84 | Elevation | Country |
|---|---|---|---|---|
| AMIG | -83.714 | 9.547 | 2778 | Costa Rica |
| BOQU | -83.692 | 9.484 | 1993 | Costa Rica |
| CUSI | -83.628 | 9.330 | 675 | Costa Rica |
| GIGA | -83.635 | 9.456 | 1407 | Costa Rica |
| LONG | -83.488 | 9.258 | 644 | Costa Rica |
| MILL | -83.687 | 9.561 | 2710 | Costa Rica |
| NIMB | -83.741 | 9.565 | 3107 | Costa Rica |
| NUBE | -83.597 | 9.388 | 1219 | Costa Rica |
| QUEB | -83.685 | 9.442 | 1261 | Costa Rica |
| RIOM | -83.782 | 9.340 | 605 | Costa Rica |
| SANM | -83.701 | 9.500 | 1973 | Costa Rica |
| TOLO | -83.701 | 9.474 | 1681 | Costa Rica |
| Alaspungo | -78.631 | 0.001 | 2484 | Ecuador |
| Alaspungo_disturbed | -78.611 | -0.010 | 2787 | Ecuador |
| Verdecocha | -78.598 | -0.121 | 3227 | Ecuador |
| Yanacocha | -78.590 | -0.121 | 3406 | Ecuador |
| Yanacocha_disturbed | -78.586 | -0.100 | 3249 | Ecuador |
